# Widespread selection relaxation in aquatic mammals

**DOI:** 10.1101/2024.09.18.613479

**Authors:** B.M. Farina, T. Latrille, N. Salamin, D. Silvestro, S. Faurby

**Affiliations:** Department of Biology, University of Fribourg, Fribourg, Switzerland; Swiss Institute of Bioinformatics, Switzerland; Department of Computational Biology, University of Lausanne, Lausanne, Switzerland; Department of Biological and Environmental Sciences, University of Gothenburg, Gothenburg, Sweden; Gothenburg Global Biodiversity Centre, University of Gothenburg, Gothenburg, Sweden

**Author notes:** Shared last authorship (D. Silvestro, S. Faurby).

**Keywords:** aquatic transitions, mammals, selection, macroevolution

## Abstract

While mammals are predominantly terrestrial, several lineages within them have independently transitioned to aquatic environments and exhibit a great variety of evolutionary changes, which ultimately lead to irreversible transitions to aquatic lifestyle. Such changes should also be detectable at the molecular level and, for instance, olfactory genes have been found to evolve under reduced selection and functionality in whales. However, the presence of a molecular signature related to aquatic transitions across other genes remains unknown as is the degree to which this affects only fully aquatic groups or also semiaquatic ones. Here, we use a Bayesian framework to investigate differences in the strength of purifying selection among terrestrial, semiaquatic and fully aquatic mammals using a set of 1000 orthologous genes, while accounting for shared ancestry, and controlling for body mass and effective population size. We found that selection relaxation linked to aquatic transitions is not only occurring in olfactory genes, but also detected across 20% of the other genes analysed here. This consistent trend of genes under weaker selection is inferred in both semi- and fully aquatic mammals although the pattern is stronger in the latter. While most of the genes analysed here likely remain functional in all mammals, their evolution is inferred to be neutral in a substantial number of aquatic species, consistently with genes that might be losing their functionality. The inferred widespread relaxation of selection among genes is consistent with a macroevolutionary scenario where secondary transitions to aquatic environment eventually become irreversible.

## Introduction

Independent secondary adaptations to aquatic habitats have occurred multiple times across vertebrates over the past 250 million years (Vermeij and Motani 2018). Among them, mammals originated in the Mesozoic and several lineages subsequently transitioned from land to aquatic environments (i.e., freshwater and marine), acquiring different adaptations in the process. These include development of novel morphological traits ranging from webbed feet in some semiaquatic groups, to major morphological and physiological changes, such as hindlimb reduction, streamlined bodies, and high hypoxia tolerance in fully aquatic species (Berta 2020; Uhen 2007; Hindle 2020). Aquatic transitions have also been shown to be linked to a consistent increase in body size likely as a consequence of thermoregulation (Gearty, McClain, and Payne 2018; Farina, Faurby, and Silvestro 2023). Due to this ensemble of phenotypic and metabolic changes, several of these evolutionary transitions to aquatic habitats are thought to be irreversible (Aboitiz 1990; Farina, Faurby, and Silvestro 2023).

The evolutionary signature of the phenotypic changes in aquatic lineages can be detected at the molecular level as well (Kelley et al. 2016; Sharma et al. 2018; Yuan et al. 2021). This has been suggested primarily in the reduction of olfaction in marine mammals, where olfactory genes have been shown to evolve under weaker purifying selection compared to terrestrial species (Kishida 2021). While whales show a reduced number of functional olfactory receptors encoded in their genome and lack the vomeronasal organ (Kishida 2021), the diminished sense of smell is not limited to fully aquatic groups, but also found in several semi-aquatic mammals, such as the North American beaver, and the sea otter (Martinez et al. 2020; Beichman et al. 2019).

Species can adapt to new environments and the genetic mechanisms responsible for these changes will involve directional selection to account for the new gene function for this evolution (Lenski 2017; Nielsen 2005). In contrast, the new environment can also lead to loss of function due to new requirements on organs and physiological processes, as in the case of olfactory receptors. These changes will usually lead to positive selection on the genes involved, in the first case, or a relaxation of purifying selection in the latter. Yet, the existence of broader the genetic changes linked to aquatic transitions beyond olfactory genes remains unclear because some lineages are more studied than others and most studies focused on a small set of genes (e.g., Nery, González, and Opazo 2013; Shen et al. 2012; Toll-Riera, Laurie, and Albà 2011; Onbe et al. 2007). Similarly, it is unclear whether these changes are limited to fully aquatic species (e.g., whales and sirenians), or if they follow evolutionary trends along a gradient of aquatic adaptations as already seen for body mass (Farina, Faurby, and Silvestro 2023) and olfaction (Martinez et al. 2020; Beichman et al. 2019).

In this study, we analysed a set of curated alignments of mammalian orthologous genes (Scornavacca et al. 2019) in a Bayesian framework (Latrille, Lanore, and Lartillot, 2021) to investigate the differences in selective strength among terrestrial, semi-aquatic and fully aquatic mammals quantifyied as gene-level ratio between non-synonymous and synonymous substitution rates. This ratio is expected to be smaller than 1 in the presence of purifying selection and both positive selection and a relaxation of purifying selection can lead to larger ratios. However, since positive selection typically affects a small number of sites and acts in a short period of evolutionary time, we expect to find gene-level estimates of the non-synonymous to synonymous rate ratio to increase mainly as a consequence of relaxed selection (Zhang 2004). We hypothesized that, since most of the evolutionary history of the mammalian clade took place in terrestrial environments and all aquatic transitions are the result of secondary adaptations (Farina, Faurby, and Silvestro 2023), there should be a relaxation of selection across several genes in aquatic mammals leading to weaker purifying selection compared to terrestrial mammals and linked to a loss or reduction of function.

## Methods

### Input data

We compiled molecular data and phylogenetic trees from OrthoMaM (Scornavacca et al. 2019), a curated database for orthologous mammalian protein-coding genes. We sampled 1,000 alignments that encompass the maximum possible number of species, and their respective gene trees, as it was shown to be a better option over a fixed species trees when estimating strength of selection (Mendes and Hahn 2016; full list of alignments is provided as Supplementary table S2). In our selection, we only included alignments with at least 900 sites and containing at least five semi or fully aquatic taxa. The datasets varied in size between 157 and 190 species (median 186) of which a maximum of 28 species were semi or fully aquatic. The alignments included between 951 and 14,271 aligned sites. We additionally sampled all 29 available alignments for olfactory receptors genes (between 924 and 1,008 aligned sites; varying from 5 to 176 species included, and from 4 to 20 aquatic ones; list of alignments is provided as Supplementary table S3), as they have been extensively studied in aquatic mammals and are known to be subjected to weaker selection in whales (Liu et al. 2019; Kishida et al. 2007a; Thewissen et al. 2011). These olfactory genes thus served as a positive control for our methodology.

We classified each mammal species in our datasets into four categories describing the level of aquatic adaptations, following the classification made by Farina, Faurby, and Silvestro (2023). This approach classifies organisms based on their morphological and life history traits, such as locomotion ability, presence of webbed feet, and reproduction under water. Under this scheme terrestrial mammals (such as horses and gorilla; class: A0) are separated from semiaquatic taxa (e.g., beaver and platypus; class: A1) with some morphological adaptations to aquatic habitats, but which spend considerable amount of time on land where they still possess high mobility capacity. We then identify as class A2 mammal species that spend considerable amount of time in water, even though they can be onshore frequently, but have limited mobility on land (pinnipeds and the sea otter). Finally, we indicate with A3 fully aquatic lineages that never leave aquatic environments (whales and manatees).

We also obtained body mass information from PHYLACINE 1.2 (Faurby et al. 2018) – as it is a fundamental trait for all organisms and a proxy for multiple physiological to ecological traits (LaBarbera 1989; Peters 1983). Finally, we compiled estimates of effective population size from Wilder et al. (2023), since it can have an impact on strength of selection (Charlesworth 2009). Whereas our dataset has complete data about body mass, effective population size lacks information for 51% of the species included here (see Supplementary Information).

### Selection estimation

We estimated branch-specific strength of selection (ω) reflecting the ratio between non-synonymous and synonymous substitution rates (*dN/dS* ratio). The average ω across codons in a protein-coding gene is usually smaller than 1, indicating that purifying selection prevents the fixation of non-synonymous mutations on most of the codons of the protein. Values of ω closer to 0 indicate stronger purifying selection, while values of ω close to 1 indicate neutral molecular evolution. We inferred branch-specific ω using the *nodeomega* model implemented in the software *BayesCode* (Latrille, Lanore, and Lartillot, 2021). The method assumes that ω, mutation rate (*μ*), and life-history traits – in our case, body mass, level of aquatic adaptation (treated here as a continuous trait, from 1 to 4) and effective population size – vary along the branches of the phylogenetic tree as a Brownian process. Missing data at the tips are imputed based on the Brownian process by the Bayesian algorithm. The model estimates the variance– covariance matrix between ω and the life-history traits as well as per-branch estimates of the rate of molecular evolution and ancestral states for the traits. It also estimates the posterior probability of a positive correlation between each pair of traits, based on the frequency of a positive covariance in the posterior sample. Thus, the posterior probability will be close to 1 for strongly supported positive correlations, close to 0 for strongly supported negative correlations, and anywhere else between 0 and 1 in case of weak correlations. We ran the analyses for 3,000 iterations with the first 1,000 used as burnin. We inspected convergence of the algorithm in Tracer (Rambaut et al. 2018).

We then assessed whether the effect of aquatic transitions on ω can be isolated from confounding effects of correlated variables. To this end, we used the estimated ω for the terminal branches to compute partial correlation coefficients between the strength of selection and aquatic adaptations, while controlling for the other variables (body mass and effective population size). We also calculated the false discovery rate as the expected number of false positives divided by the total number of positive ones to account for multiple comparisons.

Finally, to assess the robustness of our results under a different model, we ran an additional set of analyses in which ω was inferred under the *nodeomega* model, but without the inclusion of any life-history trait. We subsequently performed a phylogenetic generalized least squared analysis (PGLS) to infer how ω correlates with the level of aquatic adaptation, the effective population size, and body mass. PGLS analyses were performed using the *gls* function from the *nlme* R package, using a Brownian model to describe the phylogenetic correlation structure among species (Pinheiro and Bates 2000; Pinheiro and Bates 2023)

### Distribution of ω among genes

We explored the distribution of estimated correlations between ω and aquatic adaptations to test if 1) the number of genes with a positive correlation (i.e. higher ω in aquatic vs terrestrial species) is significantly different from that with a negative correlation and 2) if the distribution of posterior probabilities of a correlation is symmetrical. First, we performed a binomial test to assess if the number of genes with a positive covariance and correlation coefficient significantly exceeds those with negative values, under the assumption that, if aquatic adaptations do not affect the strength of selection, these numbers should be similar. To this end, we analysed the distribution of 1,000 random genes and 29 olfactory genes separately and used the *binom*.*test* function from R package *stats* (R Core Team 2024).

Second, we looked at the symmetry of the distribution of estimated posterior probabilities of a positive correlation (or p-value based on PGLS). If the relationship between species-specific ω (i.e. ω estimates on the terminal branch of the gene trees) and aquatic adaptations is random among genes we should expect the posterior probabilities to be symmetrically distributed, i.e. with some genes showing probabilities close to 0 if negatively correlated, close to 1 if positively correlated or somewhere in between if no significant correlation was detected. We fitted beta distributions to verify the presence of significant asymmetry in our estimates of the posterior probability (or p-value based on PGLS) of a positive correlation between ω and aquatic adaptations. A beta distribution spans between 0 and 1 is defined by two parameters (α and β). It can take a U shape (α and β < 1), a uniform shape (α and β = 1) or a bell shape (α and β > 1) and it is symmetric when α = β and asymmetric when α ≠ β. As posterior probability values and p-value can take values of exactly 0 or 1 due to rounding error (at which a beta distribution is not defined when α and β < 1), we discretized the beta distribution into 100 equally spaced bins. We fitted the model using maximum likelihood and compared the fit of a symmetric beta distribution (where α and β are constrained to be equal) against an asymmetric one (α ≠ β.) based on AICc scores. We also used the estimated α and β parameters to quantify the expected ratio between the number of genes with significantly positive vs significantly negative correlation, here based on the cumulative density of the beta distribution in the range 0–0.025 and 0.975–1, respectively.

## Results

### Relaxation of ω in aquatic lineages

The estimated ratios between non-synonymous and synonymous sites across species and genes were all found to be smaller than 1, indicating some level of purifying selection across all datasets. Yet, we found substantial variation in the strength of purifying selection among species in several genes (Figure 1). In particular, we found that ω significantly increases with increasing aquatic adaptation in 25.3% of the genes. After accounting for false positives, with false discovery rate of 19.8% and a 0.05 threshold, positive correlation between the two variables is still present in 20.3% of the genes. In contrast, we found that only 0.5% of the genes showed the opposite pattern, that is a decrease in ω associated with increased aquatic adaptations. This percentage drops to 0 when accounting for false positives. These results reveal the existence of a weaker purifying selection in aquatic life forms (Figure 2) consistently observed in a substantial fraction of genes.

**Figure 1.**
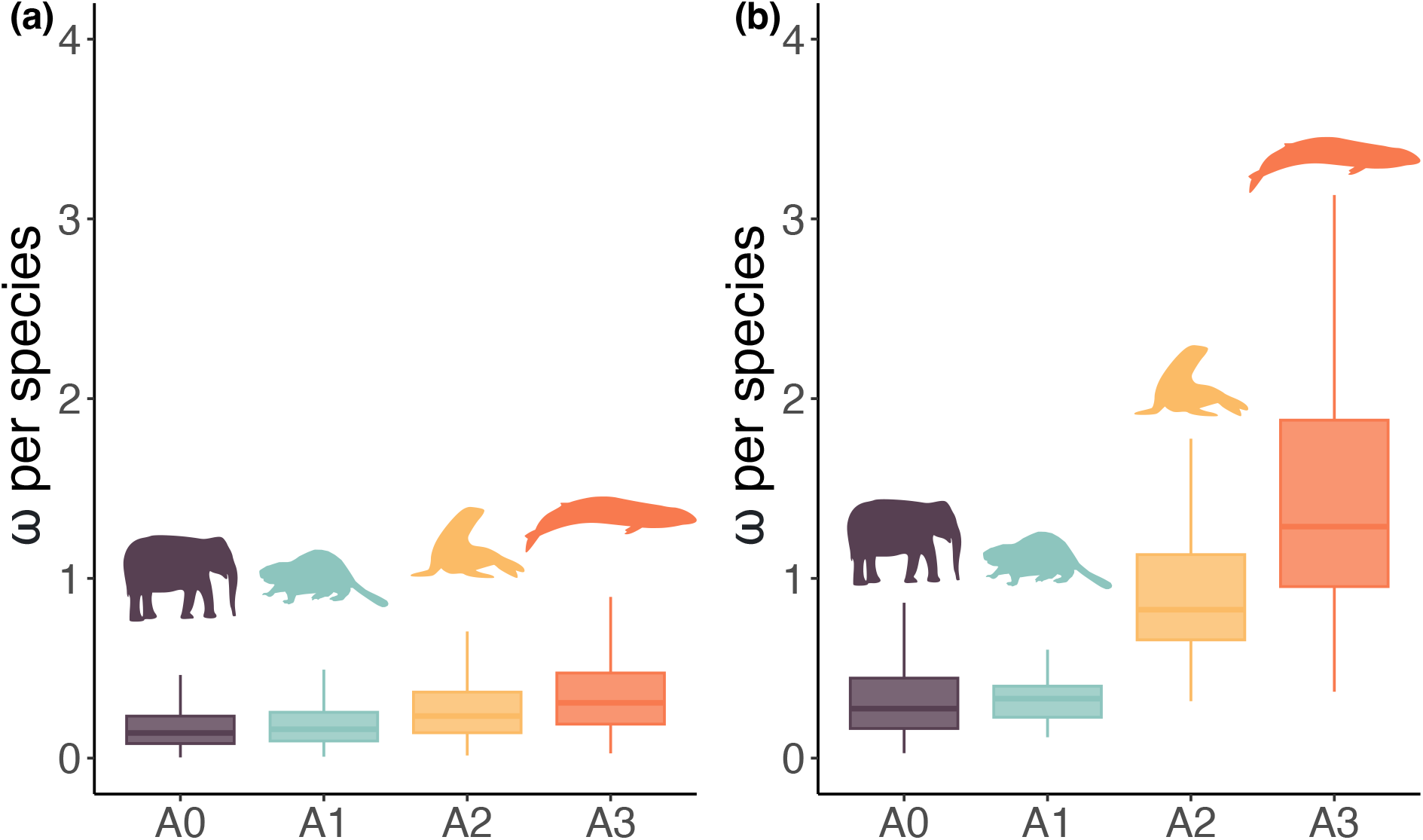
Strength of selection (ω) per species across **(a)** all miscellaneous genes per level of aquatic adaptation and **(b)** across olfactory receptors. A0 (purple) represent fully terrestrial organisms; A1 (blue green) are semiaquatic; A2 (yellow) also represent semiaquatic but those more dependent on aquatic environments; A3 (orange) are fully aquatic lineages. Coloured boxes represent interquartile ranges, and whiskers represent first quartile – 1.5 × interquartile range, and third quartile + 1.5 × interquartile range. Silhouettes from PhyloPic (Keesey, 2024) using rphylopic package (Gearty and Jones 2023).

**Figure 2.**
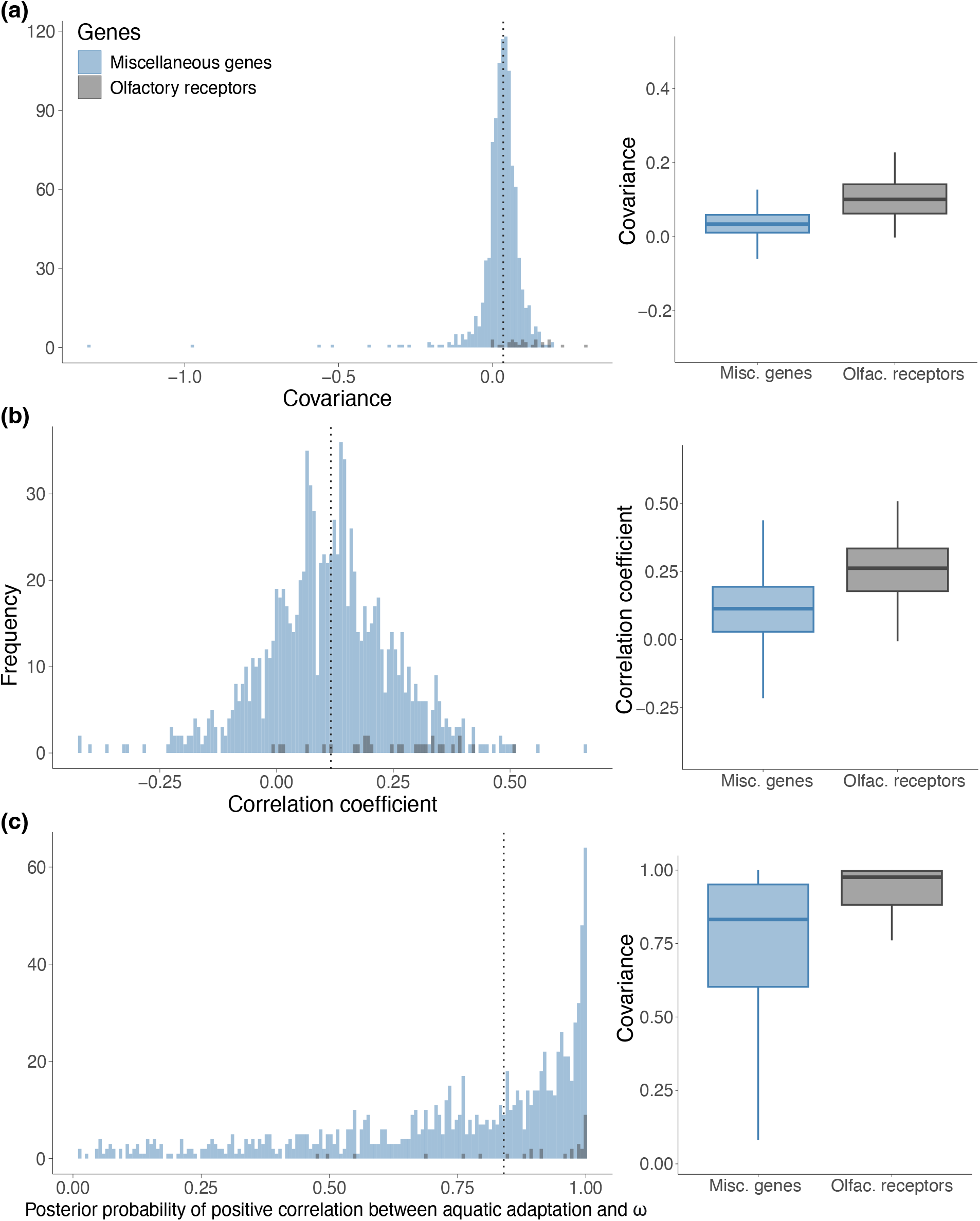
Variance–covariance matrix results for strength of selection (ω) and its association to aquatic lifestyle across all genes. **(a)** covariance **(b)** correlation coefficient and **(c)** posterior probability distributions of positive correlation between aquatic adaptations and ω; grey bars indicate olfactory receptors, whereas blue bars represent all the other genes. Dashed lines represent the median of each distribution.

The species-specific ω estimates for genes with posterior probability > 0.95 were on average 2.2 times higher in fully aquatic taxa (A3; whales and manatees) compared to terrestrials (A0), their mean estimates across genes being 0.387 and 0.175, respectively. Even in semi-aquatic groups (A2), we found a 60% higher ω (0.280 mean estimate) compared to A0, while the difference decreased to 8.8% in A1 species (0.190 mean estimate). Therefore, our results show a consistent trend with higher ω in aquatic groups, which is not restricted to fully aquatic species (Figure 1a).

Across the 253 genes with a significant positive correlation between ω and aquatic adaptations we found 74 with species-specific ω > 0.9 (therefore showing close to no evidence of purifying selection) in terrestrial species (A0), 22 in semi-aquatic (A2) and 171 in fully aquatic (A3). When accounting for the fact that many more terrestrial species are present in the dataset, we found that fully aquatic species were 25 times more likely to have ω values >0.9 compared to terrestrial and semi-aquatic species were four times more likely. Consistently with these findings, our binomial tests showed that the probability of a positive correlation coefficient is significantly higher than 0.5, i.e. the expected value under the assumption of independence between strength of selection and aquatic adaptations (Table 1, Figure 2a and b). The consistent trend toward positive correlations is also evident when fitting a beta distribution to the distribution of posterior probabilities of positive correlation between the two variables. The AICc scores strongly reject a symmetric beta distribution favouring instead a skewed distribution (Table 1, Figure 2c). This implies that relaxation in aquatic lineages (positive correlation between ω and aquatic adaptation) is 115 times more likely that the opposite pattern (Table 1).

**Table 1.** Statistical tests on the distribution of estimated correlations between ω and aquatic adaptations, and their posterior probabilities. We used a binomial test assessing the proportion of correlation coefficients across 1000 random genes and across 19 olfactory genes. and a symmetry test using beta distribution (with one and two parameters for posterior probability distribution). The probability of a positive correlation coefficient and 95% confidence interval (in parenthesis) is significantly higher than the 0.5 expected under the assumption of independence between strength of selection and aquatic adaptations. For beta distribution tests, likelihood values, estimated parameter(s), and AICc scores are provided showing that posterior probabilities were significantly values close to 1, indicating a high frequency of genes supporting higher ω in aquatic species. For the two-parameters beta distributions we also computed the probability of strong support for negative correlation and positive correlation based on the density of the beta distribution in the range 0–0.025 and 0.975–1, respectively.

|  | Miscellaneous genes | Olfactory genes |
| --- | --- | --- |
| **Corr. coef. value $\geq 0$** | Probability: 0.823 (0.798 – 0.846)<br>p-value < 0.05 | Probability: 0.966 (0.822 – 0.999)<br>p-value < 0.05 |
| **1 parameter beta distribution** | Likelihood: -4445.379<br>$\alpha$ and $\beta = 0.538$<br>AICc = 8875.022 | Likelihood: -103.238<br>$\alpha$ and $\beta = 0.216$<br>AICc = 208.624 |
| **2 parameters beta distribution** | Likelihood: -4150.588<br>$\alpha = 1.543, \beta = 0.539$<br>AICc = 8264.755<br>Prob. < 0.025 = 0.002<br>Prob. > 0.975 = 0.179 | Likelihood: -85.542<br>$\alpha = 2.460, \beta = 0.279$<br>AICc = 175.545<br>Prob. < 0.025 = 1.8e-05<br>Prob. > 0.975 = 0.485 |

[utbl]

### Other correlates

We found that the species-specific strength of selection was also significantly correlated with body mass and with estimated effective population sizes (*N*_*e*_) in a substantial number of genes (Figure S2). Specifically, 32.5% of the genes showed a significant positive correlation between ω and body mass. This fraction drops to 27.5% when accounting for false positive rate of 15.4%. In contrast, ω and *N*_*e*_ are generally negatively correlated in 29.1% of the genes, (24.1% after accounting for a 17.2% false positive rate). Increase in body size has previously been shown to be associated with aquatic lifestyle (Farina, Faurby, and Silvestro 2023), raising potential problems of co-linearity. We therefore tested if the correlations between ω and aquatic adaptations still hold after controlling for other variables (body mass, and effective population size) using partial correlation coefficients. We found that the partial correlation between ω and aquatic adaptation is still positive across a substantial fraction of genes even when controlling for the other variables (Figure 3). This shows that the correlation between ω and aquatic adaptations is not driven by associated changes in body mass and population sizes.

**Figure 3.**
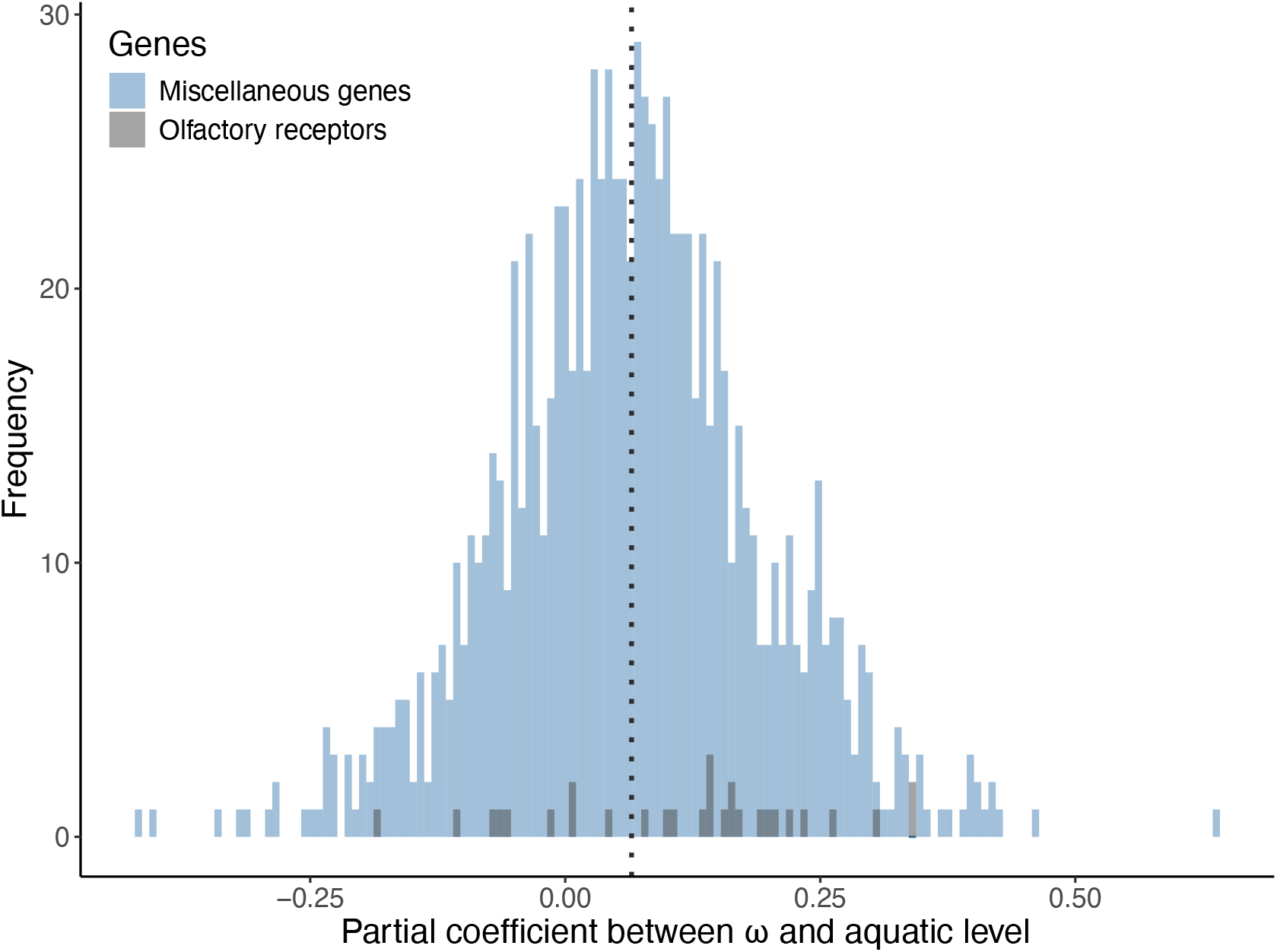
Partial coefficient between strength of selection (ω) and aquatic adaptations, after removing the effect of other correlates, namely, body mass and effective population size (Ne). Gray bars represent olfactory receptors, whereas blue bars represent other genes, and dashed line represent the median of the distribution.

### Olfactory receptors

Olfactory receptors showed a pattern of ω increase in aquatic groups (Figure 1b), as expected based on the know reduction in olfactory capabilities in several aquatic mammals. The average ω estimates for fully aquatic species was 3.6 times higher compared to terrestrial species. In semi-aquatic species (A2) the estimates were 1.7 times higher, while negligible differences were found for A1 lineages.

We found that 58.6% of the genes (53.6% correcting for false positives) are under stronger purifying selection in terrestrial lineages (not accounting for false positives; Figure 2). This confirms the known reduced olfaction ability in semi and aquatic mammals, especially marine ones. The results are also supported by the binomial and beta tests that show a trend for positive covariance and correlation coefficients, and an asymmetrical distribution of the posterior probability of positive correlation between ω and aquatic adaptations (Table 1).

### Phylogenetic patterns

A posteriori regression performed using PGLS recovered very similar patterns compared to the Bayesian analyses: covariance and correlation coefficient concerning the correlation between ω and aquatic adaptations display noncentral distributions (binomial tests with p-values < 0.05), slightly shifted to the positive side (Figure S1a and b), and p-values distribution display a clear asymmetrical beta distribution (Figure S1 and Table S1), with more genes on the left side of the distribution. PGLS recovered significant correlation in 59.6% (596 genes) of genes (false discovery rate of 7.74%). Results for the same correlation test, but for olfactory receptors, shows positive correlation in 39.8% of the genes (with 11.15% false discovery rate)

Positive correlation between ω and body mass was significant in 63.8% of the genes with false discovery rate of 7.27%. At the same time, ω and body mass for olfactory receptors show positive correlation in 15 genes (53.6%; and false discovery rate of 8.53%). Finally, 58.9% of the genes show negative correlation between ω and *N*_*e*_, while 57.0% of olfactory receptors are significant for the same correlation.

## Discussion

Evolutionary transitions toward aquatic lifestyle are followed by several phenotypic and physiological changes that have emerged independently multiple times throughout the evolution of mammals (Pyenson, Kelley, and Parham 2014; Kelley and Pyenson 2015; Reidenberg 2007). These adaptations include consistent changes, such as body mass increases (Gearty, McClain, and Payne 2018; Farina, Faurby, and Silvestro 2023; Montgomery et al. 2013), and hindlimb reduction (Gol’din 2014; Sun et al. 2022; Gingerich 2003; Gingerich, Smith, and Simons 1990). The results presented here show that these changes also include an overall decrease of purifying selection at the molecular level, consistently found across a substantial fraction of genes (∼25% in our sample of 1,000 genes). As the model used here estimates a single ω across the gene, we cannot identify whether its increase is the result of an overall relaxation of purifying selection or positive selection acting in some sites. While both explanations are theoretically possible, we interpret gene-level increases in ω as most likely the results of a relaxation of purifying selection.

In line with these observations and interpretation of the results, we found consistently higher ω in aquatic mammals among olfactory receptors, which are known to be subject to weaker purifying selection in aquatic compared to terrestrial mammals. Our findings corroborate other studies (Liu et al. 2019; Kishida et al. 2007b; Kishida 2021; Chikina, Robinson, and Clark 2016; Martinez et al. 2020; Beichman et al. 2019) that suggested diminished sense of smell in aquatic and semi-aquatic mammals – a pattern also found in other aquatic vertebrate lineages, such as marine and freshwater turtles, and sea snakes (Kishida 2021; Vieyra 2011). While baleen whales have very reduced sense of smell due to the loss of the vomeronasal organ (an important acquisition for tetrapods to be able to transition from aquatic to terrestrial environments and detect odorant molecules; Kishida 2021; González et al. 2010), toothed whales also lost their olfactory bulb and cranial nerve I, suggesting a complete lack of olfaction in this group (Kishida and Thewissen 2012). These differences in whales smelling apparatus and olfaction capacity are linked to the filter feeding behaviour of baleen whales that their use of olfaction for food detecting (Kishida and Thewissen 2012; Kishida 2021), while toothed whales rely on their echolocation ability (Berta, Ekdale, and Cranford 2014; Moss, Ortiz, and Wahlberg 2023).. Even considering groups with higher dependency on terrestrials or shallow habitats, such as pinnipeds and sirenians – which retain great part of their olfactory repertoire – aquatic mammals still have substantially fewer olfactory receptors compared to terrestrial lineages (Liu et al. 2019).

We found that a considerable proportion of the 1000 miscellaneous genes analysed here are also under much weaker selective constraints as species become more aquatic, lending support to previous hypotheses of relaxation in some genes (Chikina, Robinson, and Clark 2016). Additionally, we found species-specific ω to be close to 1 in several aquatic species, which is an indicator of near-neutral evolution and that might lead to pseudogenization of some genes. Because relaxation can lead to eventual loss of function (see Hunt et al. 2011; Zhao et al. 2023), this weaker purifying selection can contribute to the loss of “terrestrial” traits (as for the olfactory receptors) and lead to irreversible evolutionary trajectories in (semi)aquatic groups, in line with the predictions of Dollo’s law (Farina, Faurby, and Silvestro 2023). Gene losses in mammals were found to be related to diving ability (Huelsmann et al. 2019; Foote et al. 2015), for instance. At the same time, convergent relaxation has been shown to be widespread in different aquatic mammals, particularly in sensory genes and genes related to muscle proteins, possibly related to metabolic changes and body size (Chikina, Robinson, and Clark 2016).

When including additional co-variates, we also found a positive correlation between ω and body mass, which is likely due to the fact that body mass is not only linked to aquatic lifestyle per se, as an adaptation to thermoregulate underwater (Farina, Faurby, and Silvestro 2023; Torres-Romero, Morales-Castilla, and Olalla-Tárraga 2016; Adamczak et al. 2020; Goldbogen 2018), but it is also related to substitution rates, population sizes and generation time, among several other life-history traits (Martin and Palumbi 1993). Furthermore, we found that larger effective population sizes exhibit smaller ω. This is expected since relaxation can be a product of genetic drift increase, which in turn is a result of effective population size reduction (Wertheim et al. 2015). The effect of effective population size may also be linked to the effect of body size as there is a tight link between body size and population density but no strong association between range size and body size. While singling out the individual effect of aquatic transitions on selection strength from multiple co-varying factors remains difficult, our partial correlation tests show that the trend of increasing ω with aquatic adaptations holds even after controlling for changes in body mass and effective population size. The use of branch-site models of positive selection (e.g., Yang and dos Reis 2011) could help identifying specific cases in which evidence of diversifying selection might be detected in relation to aquatic transitions.

Our results show that a consistent increase in ω is not exclusive to fully aquatic taxa, like the cetaceans, but also found in semi-aquatic mammals such as seals and the sea otter. Semiaquatic adaptations are often considered intermediate between terrestrial and fully aquatic ones, as water colonization probably occurred in a stepwise manner (Farina, Faurby, and Silvestro 2023). This finding extends previous observations that amphibious groups display diminished olfactory capacity (Martinez et al., 2020; Vieyra, 2011) and convergent gene losses (Huelsmann et al. 2019), to a broader set of other genes.

## Conclusion

Mammals exhibit a trend of decreased purifying selection as species become more aquatic across large parts of their genome, not only olfactory receptors or sensory related genes. This finding adds to the growing evidence that aquatic adaptations are detectable at the molecular level. While the genes analysed here are likely still functional in most species, our analyses revealed that in several instances they have been evolving under near-neutral selection in semi and fully aquatic mammals, possibly contributing to the irreversibility of their adaptations to aquatic settings. At the same time, strong selection in terrestrial realms highlight the importance of these genes for life on land, playing the fundamental role of maintain these functions in terrestrial groups.

## Funding

BMF received funding from the Swiss Government Excellence Scholarship (2021.0350 to BMF). SF received funding from the Swedish Research Council (VR: 2021-04690). DS received funding from the Swiss National Science Foundation (PCEFP3_187012), from the Swedish Research Council (VR: 2019-04739), and from the Foundation for Environmental Strategic Research, Sweden (BIOPATH).

## Data and code availability

All the data and codes associated with this study will be made available after peer review.

## Notes

### Competing Interest Statement

The authors have declared no competing interest.

